# Draft genomes of the fungal pathogen *Phellinus noxius* in Hong Kong

**DOI:** 10.1101/209312

**Authors:** Karen Sze Wing Tsang, Regent Yau Ching Lam, Hoi Shan Kwan

## Abstract

The fungal pathogen *Phellinus noxius* is the underlying cause of brown root rot, a disease with causing tree mortality globally, causing extensive damage in urban areas and crop plants. This disease currently has no cure, and despite the global epidemic, little is known about the pathogenesis and virulence of this pathogen.

Using Ion Torrent PGM, Illumina MiSeq and PacBio RSII sequencing platforms with various genome assembly methods, we produced the draft genome sequences of four *P. noxius* strains isolated from infected trees in Hong Kong to further understand the pathogen and identify the mechanisms behind the aggressive nature and virulence of this fungus. The resulting genomes ranged from 30.8Mb to 31.8Mb in size, and of the four sequences, the YTM97 strain was chosen to produce a high-quality Hong Kong strain genome sequence, resulting in a 31Mb final assembly with 457 scaffolds, an N50 length of 275,889 bp and 96.2% genome completeness. RNA-seq of YTM97 using Illumina HiSeq400 was performed for improved gene prediction. AUGUSTUS and Genemark-ES prediction programs predicted 9,887 protein-coding genes which were annotated using GO and Pfam databases. The encoded carbohydrate active enzymes revealed large numbers of lignolytic enzymes present, comparable to those of other white-rot plant pathogens. In addition, *P. noxius* also possessed larger numbers of cellulose, xylan and hemicellulose degrading enzymes than other plant pathogens. Searches for virulence genes was also performed using PHI-Base and DFVF databases revealing a host of virulence-related genes and effectors. The combination of non-specific host range, unique carbohydrate active enzyme profile and large amount of putative virulence genes could explain the reasons behind the aggressive nature and increased virulence of this plant pathogen.

The draft genome sequences presented here will provide references for strains found in Hong Kong. Together with emerging research, this information could be used for genetic diversity and epidemiology research on a global scale as well as expediting our efforts towards discovering the mechanisms of pathogenicity of this devastating pathogen.

## Background

*Phellinus noxius* is a pathogenic soil-dwelling white-rot basidiomycete fungus belonging to the Hymenochaetaceae family. It causes the devastating Brown Root Rot (BRR) disease prevalent in tropical and subtropical countries and has a broad host range of over 260 species globally [1]. This fungal pathogen has become a growing concern in many countries in recent years as it would lead to swift deterioration of the health of the plant host within a year if left untreated. The pathogen is extremely infectious, and can be spread easily over short distances through root-to-root contact or contact with infected soil or wood pieces [2, 3], or through air-borne basidiospores for long distance dispersal [4]. It targets the roots and water transport system of the host plants, causing root mortality and compromising tree stability.

The pathogen has the ability to survive in decayed root tissue in soil for over 10 years, whilst its basidiospores can survive up to 4.5 months in soil, making it difficult to control and eradicate [5]. The aggressive nature and high infectivity of *P. noxius* have made it a global concern as it is particularly prevalent in urban settings [6–12]. *P. noxius* also has a large impact on the production of commercial crop plants such the avocados in Australia [13] and plantation conifers in Japan [10]. In Hong Kong, occurrence of *P. noxius* infections has increased exponentially over the past several years, and has become subject to public scrutiny after several collapse incidents of infected trees injuring members of the public [14–15]. Although there are several currently available methods for controlling the pathogen [16–17], there is no established cure to date.

In this study, we describe draft genome sequences of four *P. noxius* strains isolated in Hong Kong to facilitate understanding of the pathogen by investigating key factors behind its virulence and pathogenicity. Isolate YTM97 was sequenced using Ion Torrent PGM, Illumina MiSeq and PacBio RSII and hybrid assembled to generate a high quality genome sequence. Illumina HiSeq reads of the same strain were used to assemble the transcriptome to improve genome annotation. The other three strains were assembled using 3x200 bp paired-end Illumina MiSeq reads using trusted scaffolds from the previous assembly. *P. noxius* genomes generated in this study represent references for local strains in Hong Kong and allow further comparisons to global strains.

## Methods

### Isolation of fungal strains

*P. noxius* strains YTM97, YTM65, SSP14, and S39 were isolated from roots of infected trees in Hong Kong (Table S1) on 2% malt extract agar amended with streptomycin, gallic acid, benomyl, and dichloran [18] and incubated at 28°C for 10 days. Pure isolates were then cultured and maintained on 2% potato dextrose agar (PDA) before DNA extraction.

### Genome sequencing and assembly

Isolate YTM97 was sequenced using Ion Torrent PGM, Illumina MiSeq (2x300 bp paired-ends) and PacBio RSII (four PacBio SMRT cells with an insert size of 20kb). The other three *P. noxius* strains were sequenced with the Illumina MiSeq platform. Illumina and PacBio sequencing was performed at the Beijing Genome Institute (BGI) of Hong Kong and Ion Torrent sequencing was performed at the Core Facilities of The Chinese University of Hong Kong.

For the isolates YTM65, SSP14 and S39, *de novo* assembly of the Illumina MiSeq raw reads was produced using the Newbler Assembler v2.8 [19]. For YTM97, FLASh v1.2.11 [20] was used to correct the Illumina reads, which were then used to correct the PacBio reads using the program proovread [21] to offset the high sequencing error rate of PacBio[1]. SPAdes v3.9.1 [22] was used for the hybrid assembly. For the other three isolates, raw Illumina MiSeq reads were *de novo* assembled using the Newbler Assembler v2.8 [19]. The quality and completeness of the assembly was assessed using the Benchmarking Universal Single-Copy Orthologs [BUSCO] v2 software [23] and assembly statistics were calculated with QUAST v4.5 [24].

### Transcriptome sequencing and assembly

RNA-seq was performed on the YTM97 strain with the Illumina HiSeq4000 (PE101) platform at BGI Hong Kong. The isolate was grown on PDA for 10 days before total RNA extraction using the RNeasy Mini Kit (Qiagen). Raw RNA reads were filtered for quality and the remaining high quality reads were mapped against the genome assembly using Tophat 2.0.14 [25] and assembled into transcripts using Cufflinks 2.2.1 [26].

### Genome annotation

Genome annotation was performed *de novo* using GeneMark-ES [27] and AUGUSTUS v3.2.3 [28]. Predicted proteins were annotated using BLASTp and imported into Blast2GO v4.1 [29] for gene ontology (GO) assignment. Eukaryotic orthologous groups (KOGs) were assigned by RPSBLAST v2.2.15 using the KOG database [30] on the WebMGA server [31]. Carbohydrate active enzymes (CAZymes) were predicted using family classification from the CAZy database on the dbCAN web server [32] and compared to those in nine other white rot fungi. Orthologous gene clusters were assigned using OrthoVenn [33]. RepeatMasker v4.0.7 [34] was used to find repeats in the assembled genome using cross_match [35] for hits. Protein-based RepeatMasking [34] was also used to provide a second reference for known repeats and RepeatScout [36] was used to predict *de novo* unknown repeats. Secreted proteins were identified using SignalP v4.1 [37] and ProtComp v.9.0 [38]. Putative virulence factors were predicted using the Database of Fungal Virulence Factors (DFVF) [39] and Pathogen Host Interaction Database (PHI-base) [40] using a blast cutoff value of 1e^−10^.

### Phylogenetic analysis

The phylogenetic placement of *P. noxius* was inferred using LSU sequences of closely related members in the Hymenochaetaceae family retrieved from GenBank [Larsson et al 2006]. The sequences were clustered using ClustalW [41] and quality-filtered using Gblocks v0.91 [42]. Maximum composite likelihood trees were constructed using MEGA v6.06 [43] based on the Kimura 2-paramater model.

Statistical confidences on the inferred relationships were assessed by 1000 bootstrap replicates. *Oxysporus corticola, Exidiopsis calcea* and *Protodontia piceicola* were included as outgroups.

## Results and discussion

### A high-quality genome of *P. noxius* from Hong Kong

Four *P. noxius* strains were isolated from infected trees around Hong Kong (Table S1). The YTM97 strain was sequenced using Ion Torrent PGM, Illumina MiSeq and PacBio RSII. Hybrid and non-hybrid *de novo* genome assemblies were performed. The PacBio RSII and Illumina MiSeq hybrid genome assembly provided the most complete assembly with the least amount of contigs. The final assembly yielded an estimated genome size of 31,684,099 bp with an N50 value of 275,889 bp (Table 1). The GC content was 41.58% and there were over 457 contigs. The coverage was estimated to be approximately 40 fold.

**Table 1.**
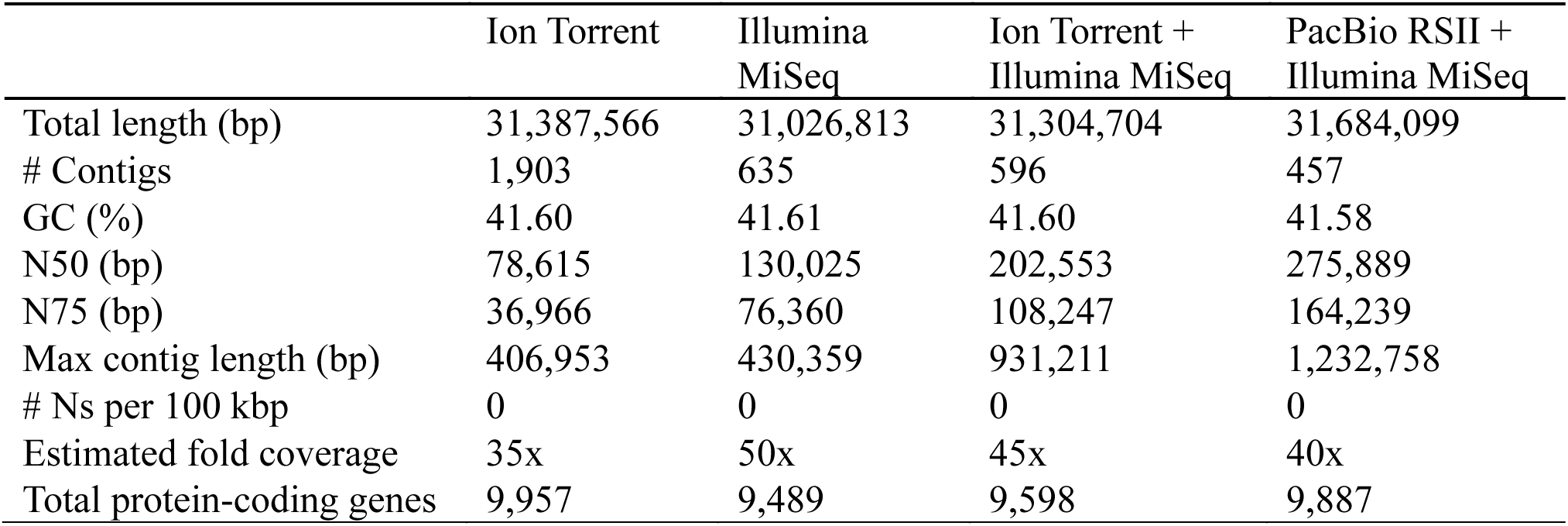
Genome assembly statistics for *P. noxius* YTM97

Illumina HiSeq4000 (PE101) RNA-seq of YTM97 yielded 74,968,996 raw reads, with a Q20 score of 98.31%. The assembled transcriptome was used to predict a total of 9,887 protein-coding genes. Genome assembly and annotation comprehensiveness recovered 279 complete BUSCOs (96.2%) suggesting the assembly is of high quality (Table S2).

The other three *P. noxius* strains were sequenced using Illumina MiSeq (2x300 bp paired end) and assembled using Newbler v2.8. The final assembly sizes varied between 30.8 – 31.9 Mb, encoding between 10,911 – 11,101 protein-coding genes (Table 2). N50 values and estimated genome completeness of these assemblies were considerably lower than those of the high quality assembly of YTM97, indicating that these assemblies were only partially complete. A pan-genome analysis for orthologous gene clusters using OrthoVenn for the *P. noxius* strains indicated that the core genome consisted of 6,744 orthologous gene clusters, whilst the pan-genome consisted of 9,399 orthologous gene clusters (Figure 1).

**Table 2.** Genome assembly statistics for all four *P. noxius* strains

| Strain | YTM97 | YTM65 | SSP14 | S39 |
| --- | --- | --- | --- | --- |
| Total length (bp) | 31,684,099 | 30,898,337 | 31,989,945 | 31,622,185 |
| # Contigs | 457 | 5,240 | 4,719 | 4,466 |
| GC (%) | 41.58 | 41.52 | 41.51 | 41.50 |
| N50 (bp) | 275,889 | 8,236 | 10,463 | 11,000 |
| N75 (bp) | 164,239 | 4,285 | 5,225 | 5,522 |
| Max contig length (bp) | 1,232,758 | 165,448 | 157,720 | 160,989 |
| # Ns per 100 kbp | 0 | 0 | 0 | 0 |
| Estimated fold coverage | 40x | 50x | 46x | 48x |
| Total protein-coding genes | 9,887 | 10,911 | 11,101 | 10,962 |
| Estimated genome completeness | 96.2 % | 69.6% | 81.1% | 75.8% |

**Figure 1.**
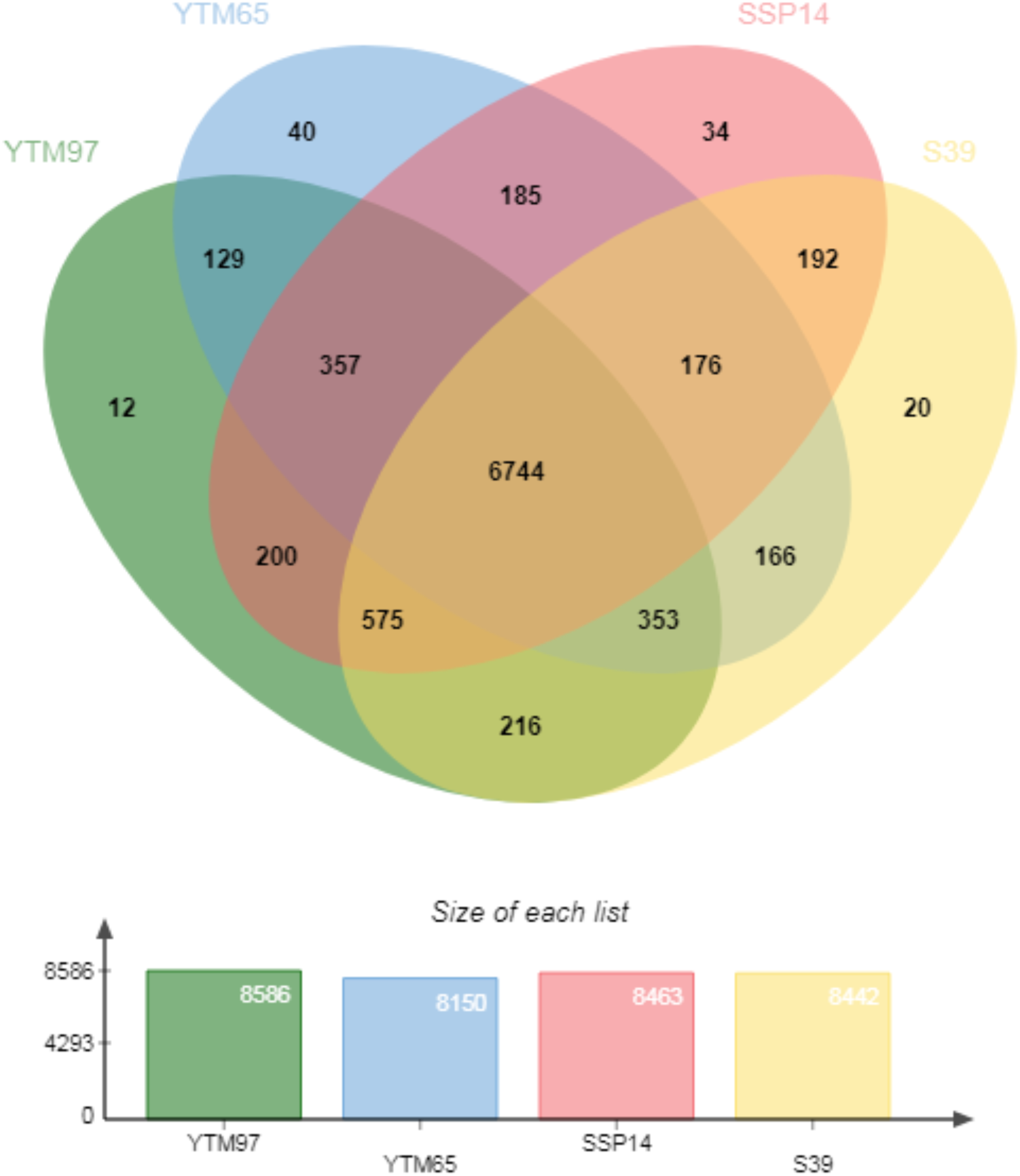
The pan genome orthologous gene clusters for the four Hong Kong strains of *Phellinus noxius*.

### Genomic features

Out of the 9,887 predicted protein-coding genes, at least one GO term was assigned to 8,690 genes (xx%). The most represented functional categories were primary and cellular metabolic processes, mainly located intracellularly (Figure S1). KOG analysis revealed that the majority of proteins were assigned to general function, signal transduction, posttranslational modification, protein turnover, transcription, as well as intracellular trafficking, secretion and vesicular transport (Figure S2). The distributions of CDS regions, transposable elements, secreted proteins, and GC skew of the 50 largest contigs, representing 21 Mb of the final 31 Mb assembly, of the high-quality genome were visualised using Circa (Figure 2). SignalP and ProtComp were used to identify 290 potential secreted signal peptides using stringent parameters that eliminated peptides with more than one transmembrane domain and glycosylphosphatidylinositol anchors. The majority of these peptides (48%) were found to be secreted extracellularly, consistent with the degradative nature of this pathogen. The distribution of these secreted proteins was uneven in the genome (Figure 2), with large stretches of contigs containing no secreted protein-coding genes.

**Figure 2:**
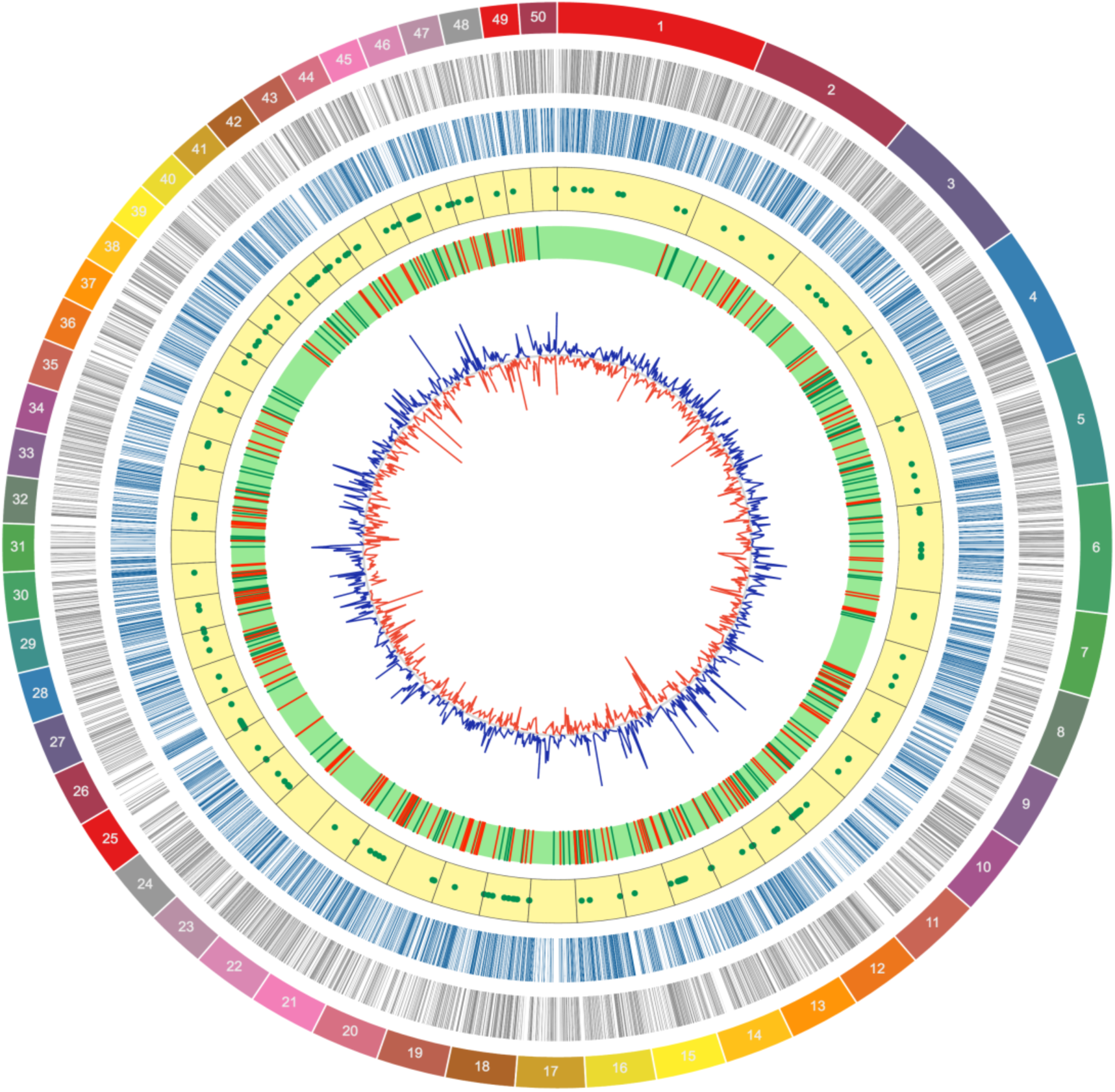
Circular representation of the genome assembly with the 50 largest contigs (21Mb) out of a complete 31Mb assembly. From outside to inside –1: Sizes of the 50 largest contigs; 2: CDS forward strand; 3: CDS reverse strand; 4: Transposable elements (TEs); 5: Secreted proteins; green – forward strand, red – reverse strand; 6: GC skew (10 kb sliding window).

### Repeats and transposable elements (TE)

RepeatScout was used for *de novo* repeat detection before running through RepeatMasker for improved accuracy and identified a total of 1,077,456 bp of repetitive elements which constituted for 3.38% of the whole genome (Table S3). The majority of the repeats belonged to simple repeats (1.62%) and LTR elements (1.26%), with only five elements showing no homology to any existing repetitive sequences. The distribution of transposable elements in the *P. noxius* genome was uneven (Figure 2), with some contigs containing only one element. The retrotransposon content in Basidiomycota genomes ranges between 3,108 copies in *Postia placenta* [44] to 47 copies in *Cryptococcus neoformans* [45]. A total of 546 LTR elements were detected in *P. noxius*, which is slightly lower than the average of 607 found in Basidiomycota [46]. However, in regard to the density per Mb, *P. noxius* has one of the highest LTR ratios due to its compact genome (Table S4). The majority of LTRs in *P. noxius* belonged to the Gypsy/DIRS1 category.

### Low levels of local and global genetic diversity of *P. noxius*

The phylogenetic relationships between *P. noxius* isolates and its closest relatives were determined using the maximum composite likelihood method with a bootstrap consensus inferred from 1000 replicates. The analysis included LSU sequences deposited in GenBank from 46 members of the Hymenochaetaceae family, with *Oxysporus corticola, Exidiopsis calcea* and *Protodontia piceicola* as outgroups. The resulting phylogenetic tree (Figure 3 and Table S5)showed well-defined clustering of different genera in concordant to previous findings [47], with the exception of the *Phellinus* genus, which formed multiple separate clades scattered throughout the tree. This indicates that the classification of the *Phellinus* genus may warrant revision or at least reconsideration. *P. noxius* clustered with the rest of the Hymenochaetaceae family with a 98% bootstrap value, but formed its own clade separated from the other members of the *Phellinus* genus.

**Figure 3:**
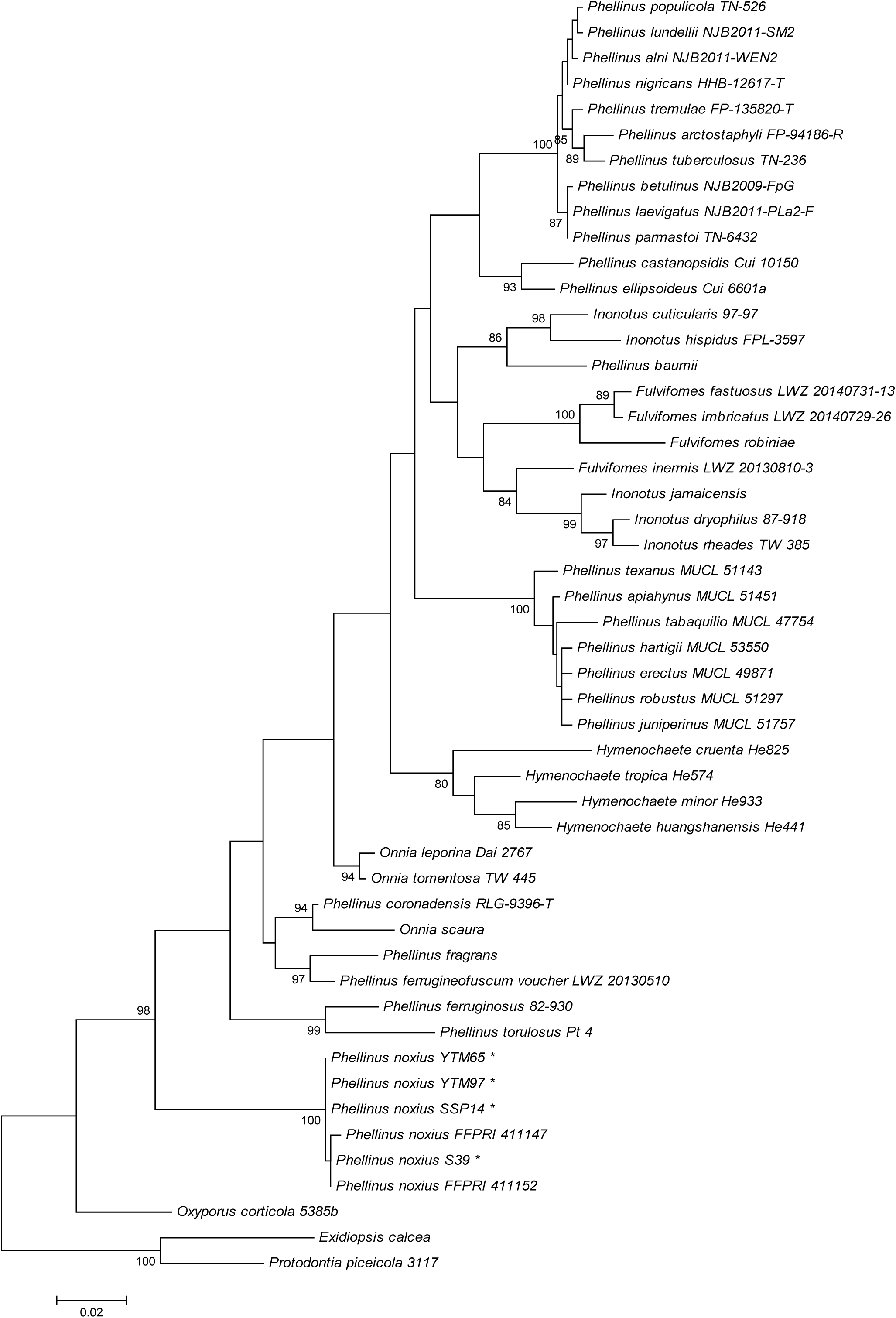
Maximum composite likelihood phylogenetic analysis using LSU sequences of *P. noxius* strains and 46 members of the Hymenochaetaceae family with a bootstrap consensus inferred from 1000 replicates.

To identify the level of genetic diversity among the Hong Kong *P. noxius* strains, single-nucleotide polymorphisms (SNPs) were called by mapping the reads of the three strains to the reference YTM97 strain. Strains YTM65, SSP14 and S39 showed similar levels of genetic variation, with 0.66, 0.63 and 0.66 SNPs per kb, respectively (Table S6). Further comparative analysis based on a total of 5,965 single-copy genes revealed the highest similarity between YTM97 and S39 (98.5%) and the lowest similarity between SSP14 and YTM65 (98.1%), suggesting that there is none or very little genetic variation between locations (Table S7). This could further suggest that *P. noxius* in these locations may have originated from the same source.

To resolve the phylogenetic relationship between *P. noxius* strains on a global scale, internal transcribed spacer (ITS) sequences from 20 strains were obtained from GenBank. Maximum composite likelihood analysis revealed that the majority of strains formed one clade (Figure 4 and Table S8) with 99% similarity, suggesting that the pathogen has little genetic divergence on a local and global scale. Most strains from the same countries were grouped together with the exception of those from Singapore, Gabon, Malaysia and Australia, suggesting that strains from these countries do not originate from the same sources.

**Figure 4:**
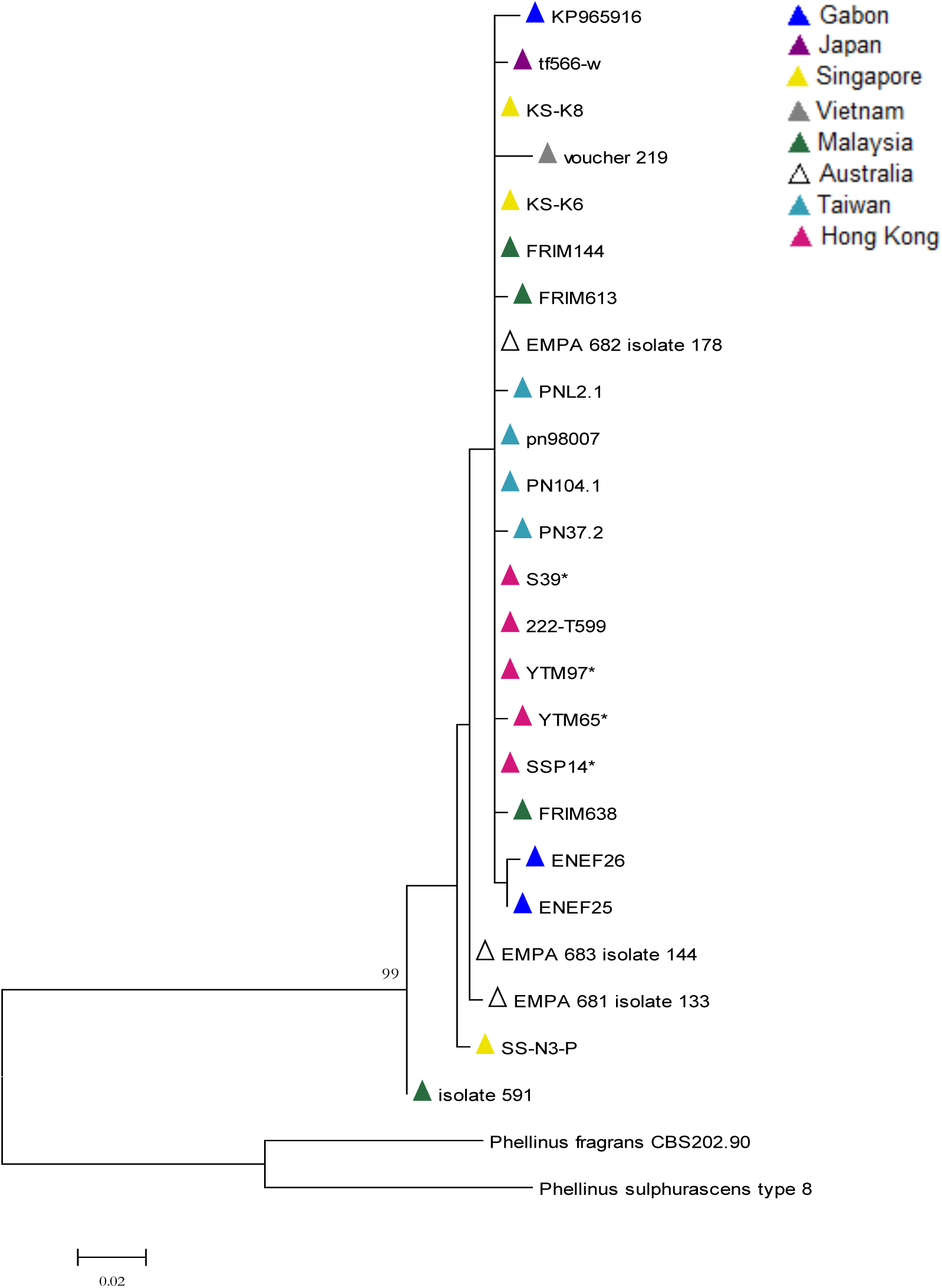
Maximum likelihood phylogenetic analysis using the ITS sequences of *P. noxius* strains from different countries with a bootstrap consensus inferred from 1000 replicates.

### CAZyme analysis

CAZymes play a large role in the pathogenicity and virulence of white-rot pathogens so the degradation capabilities of *P. noxius* were compared against nine known white-rot fungi here. Basidiomycete white-rot fungi were used in this comparison, with the exception of *Fusarium graminea.* The plant pathogens *Phanerochaete carnosa, Stereum hirsutum, Fomitiporia mediterranea, Trametes versicolor*, and *Fusarium graminea* were included due to their similar pathogenicity to *P. noxius*.

The *P. noxius* genomes encoded 400 to 477 CAZymes with a larger number of carbohydrate esterases (CE) compared to the other fungi (Figure S3). It possessed an average number of auxiliary activities enzymes (AA), which is characteristic of white-rot fungi. However, the distribution of enzymes in each AA class differed from the other plant pathogens. YTM65, SSP14 and S39 possessed 10 to 11 copies of AA1 class enzymes, which encode for laccases, ferroxidases and laccase-like multicopper oxidases used for lignolytic activities. However, only one copy was present in YTM97 (Figure 5 and Table S9), suggesting that lignin was not a major substrate for this strain and that *P. noxius* possessed the ability to efficiently utilize other carbohydrate sources. AA4 and AA7 were both present in all *P. noxius* strains but mostly absent in other plant pathogens and white-rot fungi, suggesting an increased efficiency in processing aromatic compounds, detoxification and biotransformation of lignocellulosic compounds [48]. In addition, higher numbers of CE1, CE10, GT48, GH105, and GH109 enzymes in *P. noxius* suggest that the fungus utilizes xylans, cellulose, hemicellulose, and pectin substrates efficiently in addition to lignin. The ability to degrade a wide range of substrates revealed by the CAZyme profiles and the non-specific host range of *P. noxius* together might partially explain the aggressive nature and virulence of this pathogen.

**Figure 5:**
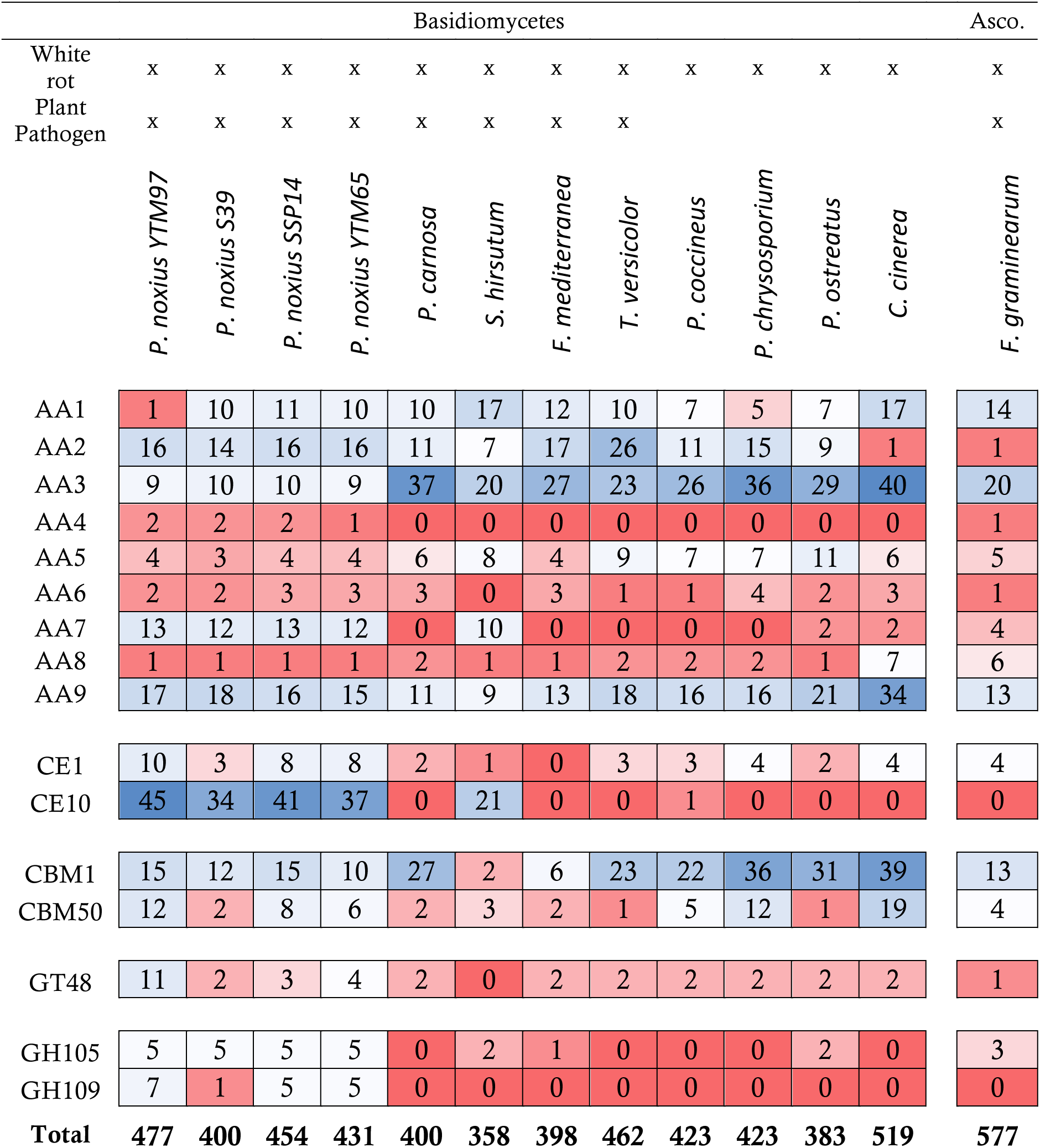
Heatmap of selected CAZymes found in *P. noxius* strains and other white rot Basidiomycetes

### Pathogenicity and virulence genes

*P. noxius* is known for its virulent nature toward a large number of hosts but to date, no studies have been performed to identify the mechanisms or genes that are involved in its pathogenesis. PHI-base is a collection of genes that have been experimentally proven to participate in pathogen and host interactions in different species. With an E-value threshold of ≤ 1e^−10^, 1,968 PHI genes were identified in YTM97. A total of 234 genes were annotated as vital for its pathogenicity, mutations or deletions of which could result in a loss of pathogenicity phenotype. Uniprot GO assignment revealed that many of these genes are involved in protein transport, autophagy, peroxisome biogenesis and the growth of appresoria and its ability to penetrate plant tissues.

Apart from virulence factors related to host interactions, a general search for putative virulence factors was performed using the DFVF with an E-value threshold of ≤ 1e^−10^. Around 3,482 putative virulence factors were predicted and transcriptional repressor Tup1, P21-rho-binding domain, DEAD/DEAH box helicase, AAA, 1,3-beta glucan synthase, cytochrome p450, cellulase, WD40, chitin synthases 1 and 2, and beta glucan synthesis-associated protein (SKN1) were in high abundance. Disruption of the *tup1* gene causes a reduction in the mating and filamentation capacity [49]; DEAD/DEAH box helicases are crucial in the signaling pathways for host and pathogen interactions [50]; beta glucan synthases and chitin synthases are vital for the biogenesis of cell walls; and P450s have now been linked to increased virulence in several pathogens [51–53]. The combination of all these virulence factors could explain the aggressive nature of *P. noxius*.

## Conclusion

The current understanding of the pathogenicity and mechanisms of infection of this pathogen are limited. Here, we present the genome sequences of four strains of *P. noxius*, the cause of the devastating brown root rot disease, posing threats to a wide range of tree hosts and crop plants globally. We were able to reveal the large repertoire of CAZymes present, possibly explaining the ability of this pathogen to kill a tree in a short amount of time and its ability to propegate and spread quickly. It has an unspecific host range and ability to degrade carbohydrates quickly, in addition to a large quantity of putative virulence factors, making it difficult to develop prevention methods.

Comparisons between the Hong Kong strains revealed the low genetic diversity present with very similar genome sequences, suggesting that these strains originated from the same source. With LSU phylogenetic analysis of the Hymenochaetaceae family, we found *P. noxius* clustered in a distinct clade away from other family members. The *Phellinus* genus is not clearly resolved and formed multiple clades throughout the phylogenetic tree, unlike other family members who formed distinct clusters within its own genera. Location based ITS analysis also revealed no significant clustering with strains from different countries, further suggesting low genetic diversity present. However, information on the Hymenochaetaceae family is limited, with only 2 fully sequenced genome sequences available making full comparative genomics difficult.

In summary, the genome sequences and information reported in this study are an important reference point for Hong Kong strains and is a valuable resource for researchers to perform genetic diversity and epidemiology research on a global scale. This information along with emerging research and the future availability of sequences from other countries could help track the spread of brown root rot disease as well as expediting our efforts towards discovering the mechanisms of pathogenicity of this devastating pathogen.

## Availability of data and materials

The datasets generated and analysed during the current study are available under BioProject ID PRJNA415882 (<u>http://www.ncbi.nlm.nih.gov/bioproject/415882</u>)

## Funding

This project was jointly funded by the Environmental and Conservation Fund and Woo Wheelock Green Fund (ECF Project 03/2014).

## Table legends

**Table S1.** Table showing locations and host species of *P. noxius* infected trees used in this study

**Table S2.** Table showing the BUSCO statistics for YTM97

**Table S3.** RepeatMasker and RepeatModeler results for YTM97

**Table S4.** LTR density ratios of *P. noxius* and select Basidiomycota

**Table S5.** GenBank accessions for Hymenochaetaceae 28S phylogenetic analysis

**Table S6.** Table showing SNP density of Hong Kong *P. noxius* strains

**Table S7.** Pairwise calculations of genetic similarity of the four Hong Kong *P. noxius* strains using 5,965 single-copy orthologous genes

**Table S8.** GenBank accessions for *Phellinus noxius* location based ITS phylogenetic analysis

## Figure legends

**Figure S1.** Graphical representation of Gene Ontology process assignment for YTM97. Showing the assignment of biological processes **(A)** and the cellular locations of these processes **(B)** using GO terms.

**Figure S2.** Graphical representation of KOG assigned classes for YTM97

**Figure S3.** Predicted CAZyme numbers in white-rot fungi

